# Processing of the SARS-CoV pp1a/ab nsp7-10 region

**DOI:** 10.1101/860049

**Authors:** Boris Krichel, Sven Falke, Rolf Hilgenfeld, Lars Redecke, Charlotte Uetrecht

## Abstract

Severe acute respiratory syndrome coronavirus (SARS-CoV) is the causative agent of a respiratory disease with a high case fatality rate. During the formation of the coronaviral replication/transcription complex (RTC), essential steps include processing of the conserved polyprotein nsp7-10 region by the main protease M^pro^ and subsequent complex formation of the released nsp’s. Here, we analyzed processing of the coronavirus nsp7-10 region using native mass spectrometry showing consumption of substrate, rise and fall of intermediate products and complexation. Importantly, there is a clear order of cleavage efficiencies, which is influenced by the polyprotein tertiary structure. Furthermore, the predominant product is an nsp7+8(2:2) hetero-tetramer with nsp8 scaffold. In conclusion, native MS, opposed to other methods, can expose the processing dynamics of viral polyproteins and the landscape of protein interactions in one set of experiments. Thereby, new insights into protein interactions, essential for generation of viral progeny, were provided, with relevance for development of antivirals.

## 1.2. Introduction

The human pathogenic potential of zoonotic infections by coronaviruses was exposed by the discovery of severe acute respiratory syndrome coronavirus (SARS-CoV) as causative agent of the SARS epidemic in 2003 [1, 2]. The replicase gene ORF 1ab occupies two thirds of the CoV single-strand (+)-sense RNA genome. Initially, ORF1ab is directly translated into either replicase polyprotein pp1a (nsp1-11) or pp1ab (nsp1-16), depending on a ribosomal (−1)-frameshift [3]. Subsequently, the polyproteins undergo proteolytic processing into 11 or 16 individual nsp’s. Eventually, the nsp’s take part in forming the RTC, a membrane-anchored, highly dynamic protein-RNA complex facilitating the replicative processes [4–7].

The main chymotrypsin-like protease (M^pro^; nsp5) facilitates processing of the C-terminal part of the polyprotein [8]. Following maturation by auto-cleavage, two protomers of M^pro^ associate into one active dimeric unit that specifically targets the cleavage sites at the nsp inter-domain junctions of nsp5-11 or nsp5-16. These sites primarily contain LQ↓S or LQ↓A residues at the positions P2, P1 and P1′, but strictly conserved is only Q at P1 [9]. Being a possible drug target, structure and function of this protease were extensively investigated [10, 11].

The timely coordinated processing of pp1a is crucial for replication [12]. This was specifically shown for the processing order of the nsp7-10 region [13]. In general, the order of processing is directly dependent on the substrate specificity of M^pro^, which has been tested in peptide-based biochemical assays [14–16]. However, peptide assays interrogate merely the substrate specificity for the isolated amino-acid sequence but disregard the polyprotein's structural layout, i.e. overall conformation or accessibility.

Upon processing of the nsp7-10 region by M^pro^, the four small proteins nsp7, nsp8, nsp9 and nsp10 are released [17]. They can form functional complexes with CoV core enzymes and thereby stimulate replication. Most importantly, a complex of nsp7+8 acts as processivity factor for nsp12, the RNA-dependent RNA-polymerase (RdRp) [18]. Particularly, various oligomeric states of nsp7+8 have been reported [19–22].

Detailed studies of nsp processing and complex formation require a method that follows these dynamic processes at a molecular level. Native mass spectrometry (MS) probes and detects protein complexes in mixtures, while keeping their non-covalent interactions intact through nano-electrospray ionization (nanoESI) [23, 24]. Furthermore, collision-induced dissociation (CID) enables to decipher the stoichiometry and topology of natively sprayed protein complexes, by dissociating non-covalent interactions *in vacuo*. Thereby, it allows to reveal the landscape of protein species from a heterogeneous sample solution and to observe dynamic processes over time.

Herein, we analyzed the processing of SARS-CoV nsp7-10 and complex formation of its sequentially released nsp’s. First, the substrate specificity of SARS-CoV M^pro^ for the cleavage sites within nsp7-10 was determined in an assay with FRET-labelled peptides. Then, we explored native MS to reveal time-resolved nsp release and complex formation upon processing of a full-length SARS-CoV nsp7-10 substrate. Specifically, the analysis via CID-MS gave insights into a preferred complex of SARS-CoV nsp7+8.

## 1.3. Results

### 1.3.1. Specific substrate efficiencies from FRET peptide assay

Initially, the substrate specificity of SARS-CoV M^pro^ for nsp7-10 inter-domain junction cleavage was studied by means of FRET peptide substrates (FPS’s). The substrates FPS7-8, FPS8-9 and FPS9-10 represented the cleavage sites nsp7-8, nsp8-9 and nsp9-10, respectively. Additionally, FPS4-5 served as a positive control analogous to the highly efficient N-terminal auto-cleavage site nsp4-5 of SARS-CoV M^pro^ (nsp5). The FPS’s encompassed 12 amino-acid residues spanning positions P6 to P6’ around the cleavage site and were labelled with a fluorophore and a quencher at their N- and C-terminus, respectively. Substrate specificities were determined by monitoring the increasing fluorescence upon M^pro^ cleavage of the FPS (Figure 1 and Table S 4). As expected, the assay showed FPS4-5 to be most efficiently cleaved amongst the FPS’s tested here, with an apparent efficiency (K_cat_/K_M_) of 0.105±0.005 μM^−1^min^−1^. Compared to FPS4-5 (100 %), substrates FPS7-8 (17.1 %) and FPS9-10 (51.8 %) were less efficiently cleaved while FPS8-9 (0.1 %) was almost non-cleavable. The absolute and relative substrate efficiencies determined here agree with results of similar experiments [14, 25]. Considering that short peptides represent merely the primary structure of the cleavage site, more native-like substrates were prepared and tested to verify these results obtained with the FPS’s.

**Figure 1:**
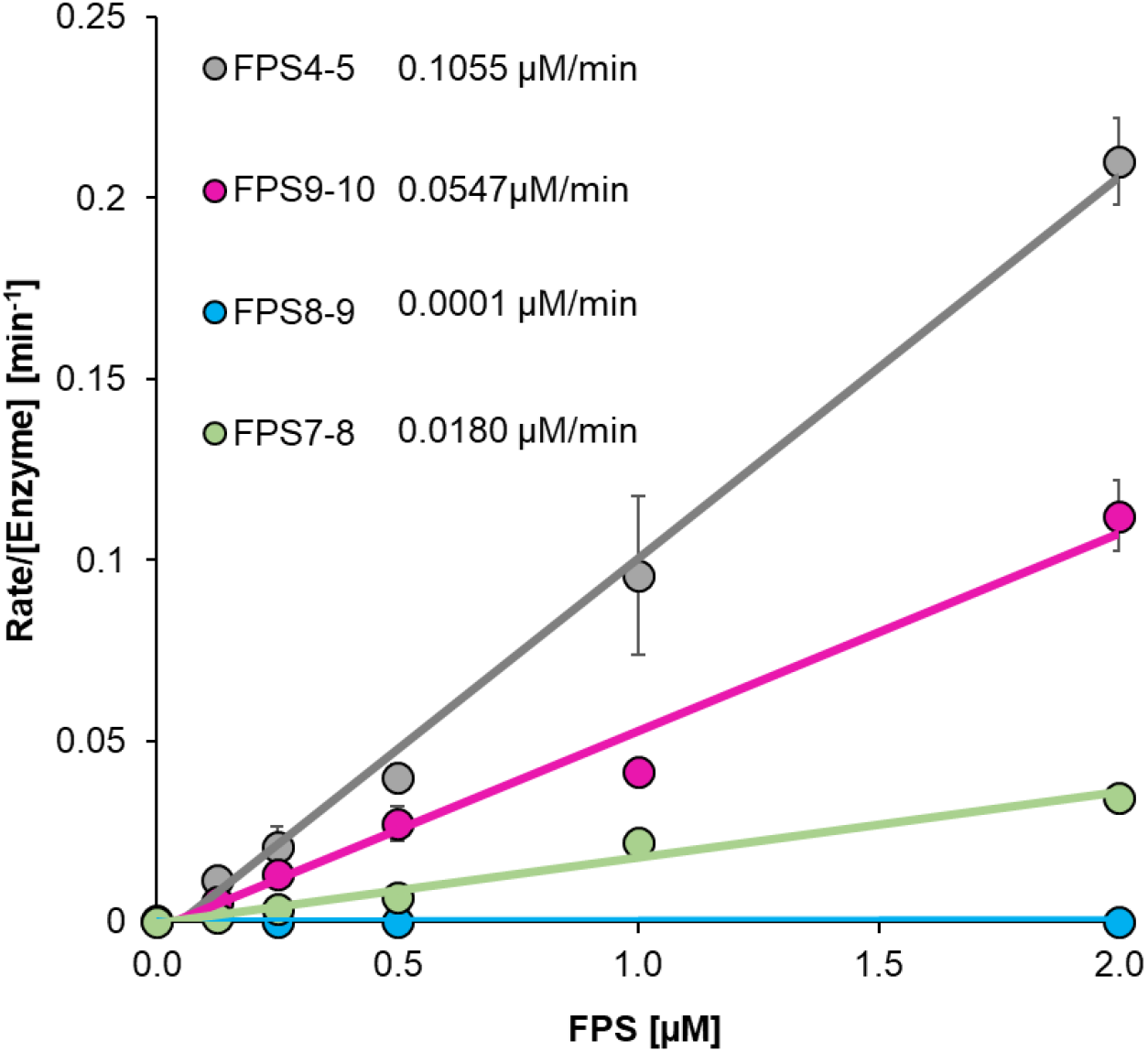
Process curves of M^pro^ (0.5 μM) with different concentrations of FRET peptide substrates (FPS) in 50 mM HEPES, 10 % Glycerol pH 7.5. The slopes correspond to the apparent k_cat_/K_M_ as given in the legend.

### Processing of nsp7-10

Next, we studied the substrate specificity of SARS-M^pro^ for folded proteins, which exhibit the structural environment of the cleavage sites beyond the amino-acid sequence. Therefore, full-length SARS-CoV His-nsp7-10 was recombinantly produced in *E.coli* and purified. This protein construct had a non-cleavable N-terminal His tag to reduce number of cleavage product species and thereby simplified interpretation of the mass spectra.

To monitor the processing dynamics, SARS-CoV His-nsp7-10 was cleaved by SARS-CoV M^pro^ in a test tube and samples were subjected to native MS. Cleavage was performed in a nano ESI compatible ammonium acetate (NH_4_OAc) solution at pH 8.0, which proved equally suitable to maintain protease activity compared to buffers with NaCl (Figure S 1). Processing products were observed as peaks in the mass spectra and assigned to protein mass species (Figure 2, Table S 3). When acquiring mass spectra at different times during processing, signals of three protein categories dominated successively, in the beginning full-length nsp7-10 substrate and M^pro^ (~1h); subsequently, intermediate products (~6 h), representing partially cleaved substrate; and in the end mature products encompassing nsp monomers and protein complexes (~20 h) (Figure S 6). Covalent and non-covalent products were distinguished via CID-MS/MS.

**Figure 2:**
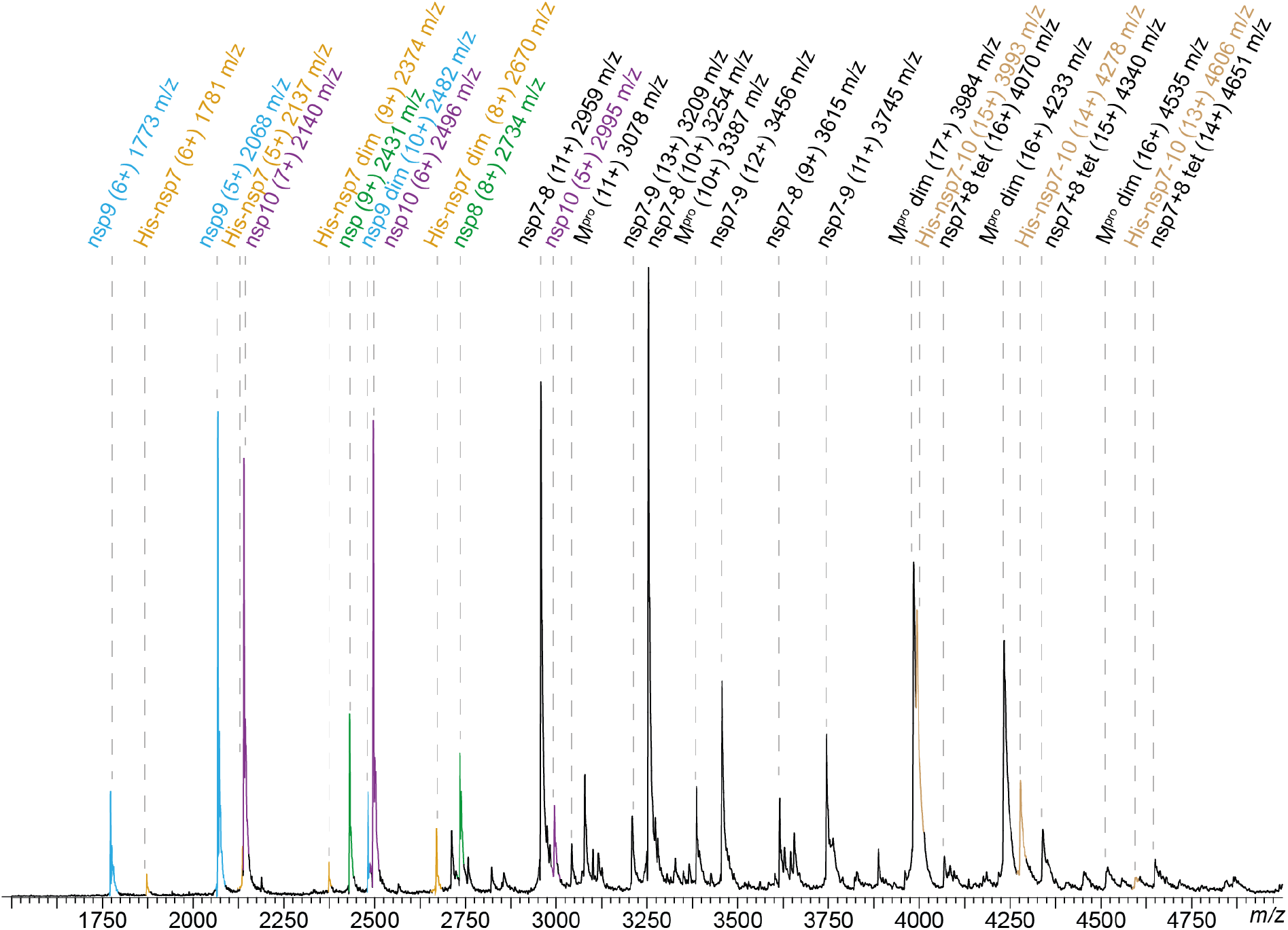
SARS nsp7-10 processing in native MS. Peak assignment in native MS after 6h. Mass species, charge states and their *m/z* are labelled. For processing, 1.25 μM SARS-CoV M^pro^ and 12.5 μM SARS-CoV His-nsp7-10 (~1:10 ratio) were incubated at 4°C in 250 mM NH_4_OAc, 1 mM DTT at pH 8.0. The components were mixed and after a defined incubation time, samples were injected into an electrospray capillary and mass spectra were acquired.

The order of reactions became evident from monitoring peak intensities over time. The relative signal intensity of M^pro^ remained stable while nsp7-10 decreased (Figure 3 A), indicating that M^pro^ depleted nsp7-10. At 20 h, no signal of nsp7-10 was detected anymore and therefore, cleavage was considered complete.

**Figure 3:**
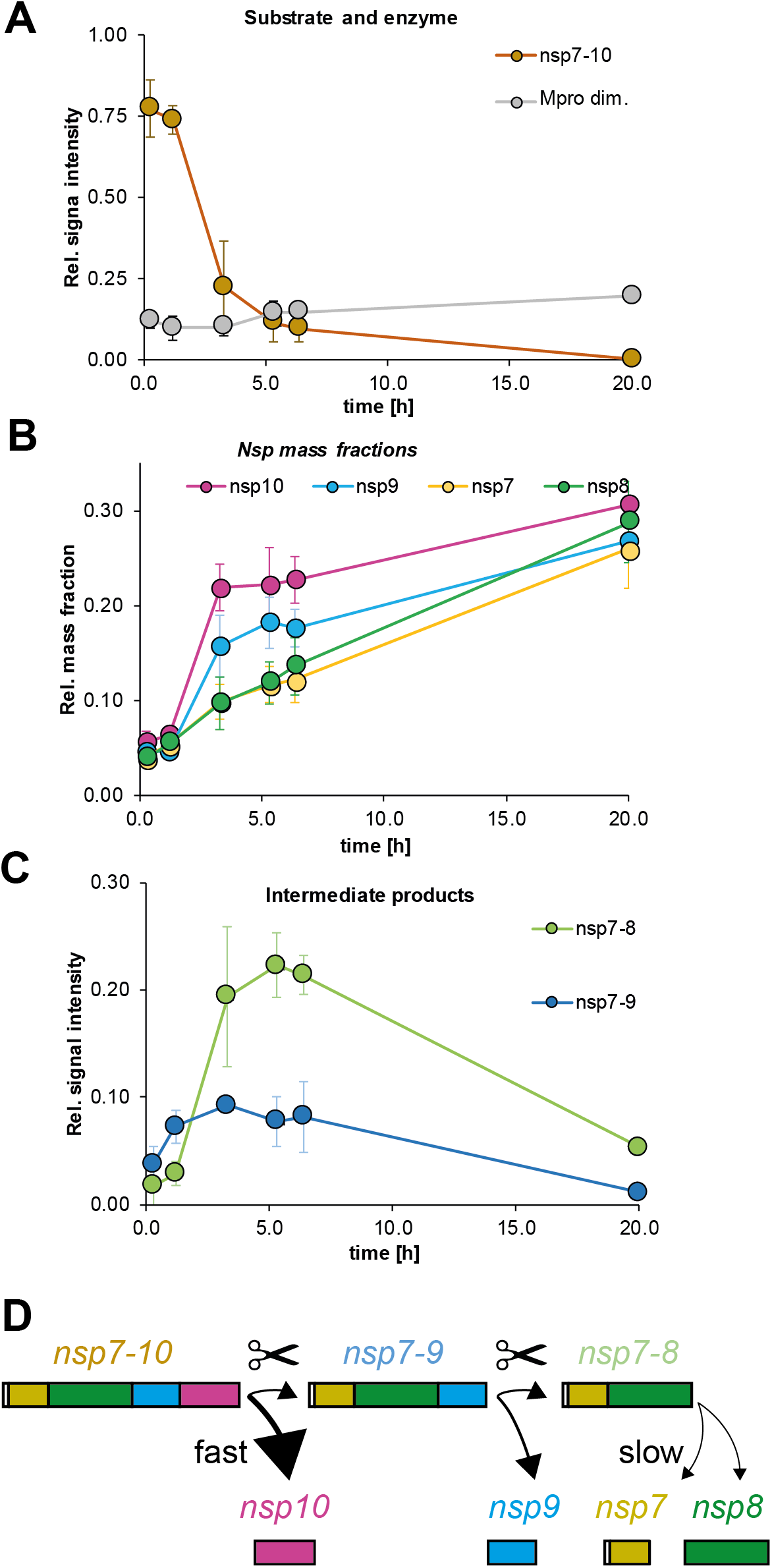
SARS-CoV nsp7-10 processing monitored by native MS: Signal over time of protease, substrate and products. **(A)** Substrate and enzyme. **(B)** Mass fractions over time show the order of nsp release. **(C)** Intermediate products. Error bars depict standard deviation (*N*=3). Time points (AVG±SD, *N*=3): 0.25±0.1 h, 1.2±0.1 h, 3.3±0.2 h, 5.3±0.3 h, 6.4±0.3 h±0.5 h. **(D)** Schematic illustration of cleavage order and efficiency.

To obtain the sequence of nsp release, the ratio of the mature nsp monomers was followed over time. However, some non-covalent interactions of the nsp’s introduced a bias to the relative peak intensities among monomers (Figure S 5). To correct for this bias, signal intensities were converted into mass fractions (MFs), which include the relative signal of each released subunit over time (Table S 2). Thereby, MFs demonstrated a processing order of the nsp9-10 site first, the nsp8-9 site second, and the nsp7-8 site third (Figure 3 B). MFs were similar at the end-point of processing (~20 h) showing that the individual nsp’s were all cleaved off nsp7-10 (Figure S 5).

To determine the relative cleavage efficiencies in the full-length substrate, we used the relative MFs between nsp10, nsp9, nsp8 and nsp7 at 3.3 h (100%, 66%, 36 % and 36 %, respectively). These values suggest decreasing cleavage efficiency from the C- to the N-terminus of SARS nsp7-10. Consistently, the cleavage pattern of the intermediate products revealed that nsp7-10 is predominantly cleaved to nsp7-9 and only then to nsp7-8 (Figure 3 C). The time-course of processing of the SARS-CoV full-length substrate as determined here via native MS corresponded well with results from SDS-PAGE using the shorter SARS-CoV His-nsp7-9 substrate (Figure 3 C, Figure S 7). Additionally, shorter substrates were also analyzed in native MS, fully in line with the results obtained SARS-CoV nsp7-10 (Figure S 8).

The results from FRET peptide assay and full-length protein cleavage are in conflict regarding the suggested cleavage order. This is particularly interesting for nsp8-9, because of its *Coronaviridae*-wide conserved NNE motif at P1’-P3’. Previously, it was suggested that this motif could confer poor substrate properties to the nsp8-9 cleavage site, which would result in a considerable influence on processing order and RTC assembly [16]. In fact, two observations made by other researchers provide indication for a peptide-specific effect: Fan et al. analyzed the secondary structure of nsp peptides and observed that nsp8-9, comprising the NNE motif and closely resembling the FPS8-9 tested here, had a higher propensity for α-helices than other nsp peptides [26, 27]. Moreover, Chuck et al. reported that β-strands in substrates promote binding to M^pro^ [28]. The results shown in our work indicate that the structural layout of the cleavage site is different when comparing peptide to full-length protein, which plays a role in endowing the nsp8-9 cleavage site with decent cleavability. Since the full-length protein possesses the structural environment of the cleavage sites, we consider results with this substrate more relevant for studying the processing pattern of the CoV polyprotein.

### 1.3.3. Protein complex formation upon processing of nsp7-10

In parallel, we applied CID-MS/MS to identify non-covalent interactions. Thereby, covalent processing intermediates were distinguished from newly formed complexes, for instance nsp7-8 intermediate and nsp7+8(1:1) dimer overlap in mass spectra (Figure S 3). Most importantly, we identified an nsp7+8 (2:2) complex, undescribed before, and subject to detailed analysis later on. Here, this complex’ intensity in native MS remained low, most likely due to interference by the non-cleavable His-tag N-terminal of nsp7. Furthermore, homo-dimers of nsp7 and nsp9 were found (Figure S 4). The MFs of nsp9 monomers and dimers (AVG±SD, *N*=3; 75±11% and 25±2%, respectively) were determined, based on the intensity ratio of the assigned peaks. Although, the X-ray structure of nsp9 was reported as a dimer, its low abundance found here is consistent with a low dimer affinity in solution [29]. So far, nsp7’s ability of dimerization in absence of other proteins has already been noted, but no function has been suggested [21] and therefore, more detailed information remains elusive. Many interactions between components of nsp7-10 have been reported in the past, however, nsp interactions other than those reported above were completely absent, despite the high sensitivity of native MS.

### 1.3.4. Complex formation of nsp7+8

Finally, the complex of nsp7+8 was investigated in detail. In order to promote interactions of nsp’s with authentic amino acid sequence, we engineered a SARS-CoV nsp7-9-His containing a cleavable His-tag at the C-terminus. Upon processing of this substrate by M^pro^, we tested the landscape of mass species via native MS. Peak envelopes were detected for the monomeric products nsp7, nsp8 and nsp9 as well as for M^pro^ monomer and dimer. While nsp9 did not interact with any of the other subunits, even after cleavage of the tag, nsp7+8 formed a hetero-tetramer of nsp7+8(2:2) (62.2 kDa) (Figure 4). This stoichiometry was predicted based on the molecular weights of the subunits (2×9.3 kDa + 2×21.8 kDa = 62.2 kDa). However, for further evidence, molecular ions of three different charge states of the putative nsp7+8 hetero-tetramer (4449 *m/z*, +14; 4153 *m/z*, +15; 2893 *m/z*, +16) (Figure 5 A) were selected as precursor and subjected to CID-MS/MS. To successively induce subunit dissociation, we increased the collision voltage (CV) stepwise. Thereby, the overall dissociation pattern was exposed (Figure 5 A and Figure S 9), which confirmed the hetero-tetrameric (2:2) complex stoichiometry.

**Figure 4:**
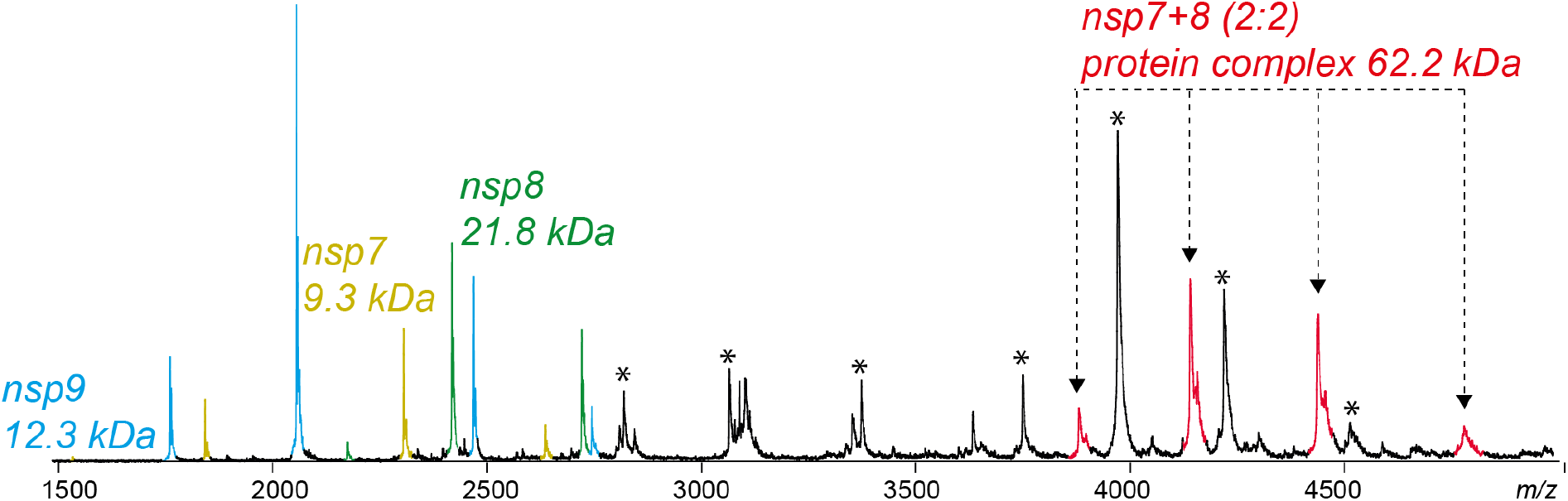
Upon cleavage of nsp7-9-His, a putative nsp7+8 (2:2) tetramer (red) emerged. Other assigned peaks correspond to M^pro^ (asterisk), nsp7 (yellow), nsp8 (green) and nsp9 (blue). For cleaving the precursor, 2 μM SARS-CoV M^pro^ was incubated with 14 μM SARS-CoV nsp7-9-His (ratio ~1:7) at 4°C in 300 mM NH_4_OAc, 1 mM DTT at pH 8.0 overnight.

**Figure 5:**
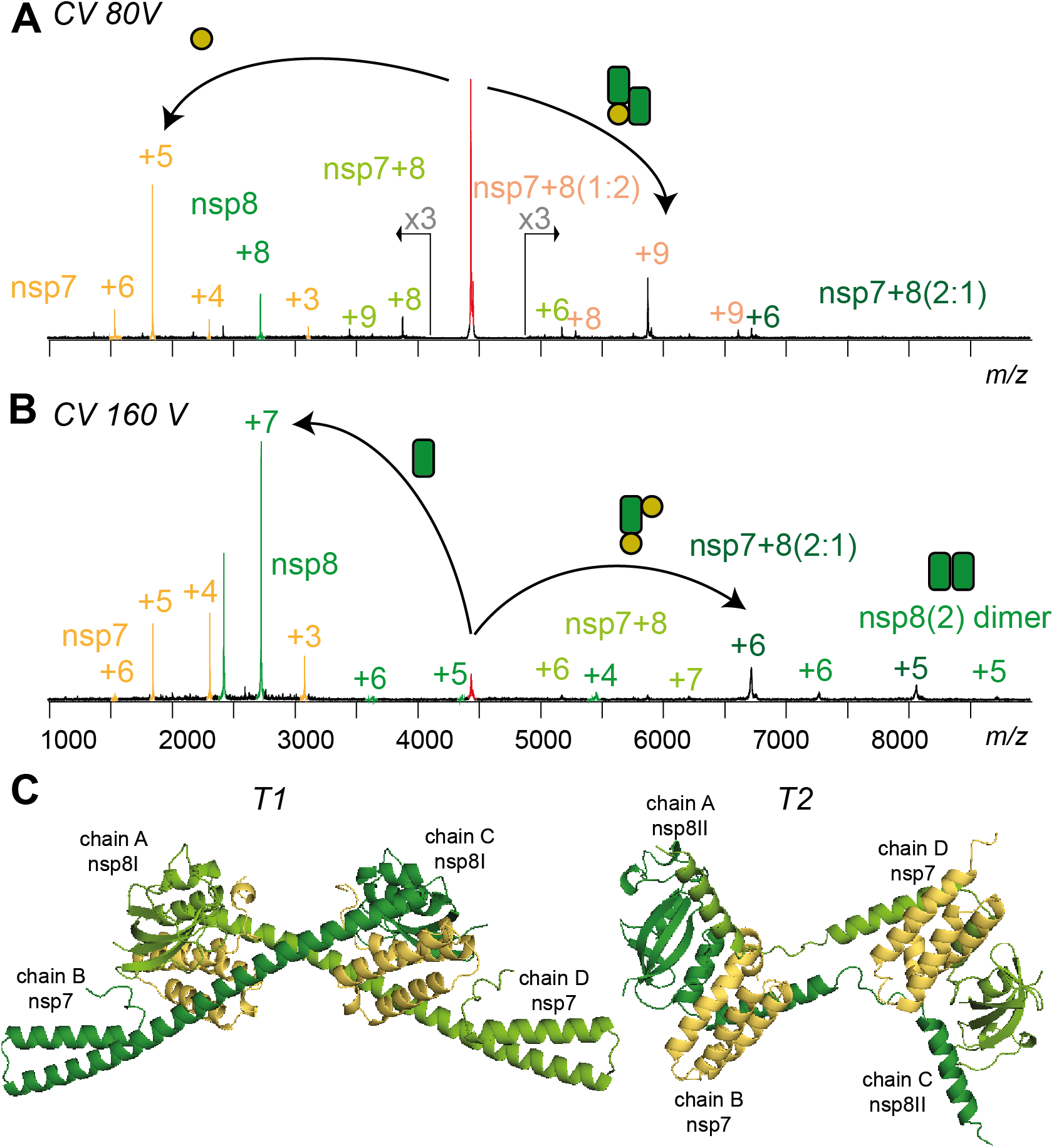
Product ion spectra showing two CID dissociation pathways of SARS-CoV nsp7+8 hetero-tetramer (+14, 4449 *m/z*) at (A) 50 V and (B) 80 V collisional energy. (C) Two conformers denoted as T1 and T2 extracted from a hexa-decameric SARS-CoV nsp7+8 (PDB 2G9T) [30] and both in line with the native MS results.

Two alternative dissociation pathways revealed the arrangement of subunits within the complex (Figure 5 B). The hetero-tetramer dissociated at lower activation energy into nsp7 and nsp7+8(1:2), and alternatively, at elevated collisional energy into nsp8 and nsp7+8(2:1). Additionally, the product ions nsp7+8(1:1) and nsp8(2) provided evidence that their subunits interacted in the complex. The alternative dissociation routes may have resulted from analogous energy absorption paths, either due to similar binding contribution of nsp7 and nsp8 in the complex or due to different gas phase conformers.

## 1.4. Discussion

Recently, the structure of the SARS-CoV RdRp complex was reported as nsp(7+8)+8+12 ((1:1):1:1)[22]. Nevertheless, the quaternary structure in the absence of other proteins could be different and an important intermediate for either stability or subsequent interactions in solution. Although, molecular structures of isolated SARS-CoV nsp7+8 have been reported as dimer (1:1) containing N-terminally chopped nsp8 and as a hexa-decamer (8:8) [20, 30]. Up to date, a hetero-tetrameric interaction has not been shown.

In terms of stoichiometry and subunit connectivity, only nsp7+8(2:2) subcomplexes of the hexadecamer resemble our results [30]. Specifically, in two conformers, T1 and T2 (Figure 5 C), nsp8 builds a scaffold with head-to-tail interaction and nsp7 subunits sandwich the scaffold without self-interaction. All interfaces are mainly hydrophobic. However, three polar bonds within or in proximity of the nsp8 shaft domain facilitate binding to nsp7. Polar bonds more readily persist in the gas phase than hydrophobic interactions, which explains higher gas phase stability for nsp7+8(1:1) over nsp8(2). The total buried surface areas of T1 and T2 are virtually equal (Table S 5). Nevertheless, the nsp8(2) dimer scaffold in T2 is compact because a loop within the N-terminal α-helix allows bending and interacting with the other nsp8, whereas in T1 the α-helix extends to interact with another tetramer in the crystal lattice. Hence, T2 represents a more likely structure of nsp7+8(2:2) in solution.

In summary, we performed a comprehensive analysis of specific processing and complex formation of a folded viral polyprotein with native MS. While it remains a challenge to perform quantitative MS from heterogeneous protein mixtures, the usage of mass fractions disentangled the mass species and allowed following the processing dynamics. Intriguingly, our results suggest that the processing pattern of SARS-CoV nsp7-10 could have evolved to tailor a remaining nsp7-8 intermediate product. This intermediate product may have independent enzymatic function [21] but finally serves as an efficient precursor for nsp7+8(2:2) hetero-tetramer formation, as stimulant of the CoV RdRp [18]. Extending our approach to processing of different CoV species and longer polyproteins will increase understanding of the formation of the functional CoV-RTC.

## 1.5. Materials and methods

### 1.5.1. Cloning and gene constructs

To generate inserts for expression plasmids for the CoV ORF1ab nsp7-10 region, DNA was amplified by PCR from commercially obtained cDNA (Eurofins scientific SE) and from DNA plasmids existing from earlier work [17, 21]. To create ends for directed cloning, the PCR products were digested by Eco31I (BsaI) (Thermo Fisher Scientific) and ligated into IBA pASK33+ and pASK35+, encoding for C- and N-terminal His-tag, respectively. The expression plasmid for SARS-CoV M^pro^ with authentic ends was created as described by Xue et al.[31].

### 1.5.2. Expression and purification

Main protease SARS-CoV M^pro^ was produced with authentic ends [31]. Briefly, the protein constructs had a GST-tag connected via an auto-cleavage site at the N-terminus and a His-tag connected via a PreScission protease cleavage site at the C-terminus. Protease was purified by affinity chromatography using Ni^2+^-NTA beads (Thermo Fisher Scientific) and dialysis into storage buffer (50 mM HEPES, 10 % glycerol pH 7.5). The His-tag was cleaved by dialysis into PreScission cleavage buffer (20 mM Tris-HCl, 150 mM NaCl, 1mM EDTA, 1 mM DTT at pH 7.0) while incubating with PreScission protease (protease to protein ratio 1:2000) (GE Healthcare) overnight. PreScission protease was removed by binding to GST-Sepharose (GE Healthcare) for 2 h and buffer exchange with centrifugal filter devices (10,000 MWCO, Amicon, Merck Millipore) into storage buffer. Purified M^pro^ was frozen in liquid nitrogen and stored at −80°C.

To produce proteins (Protein ID: P0C6X7) of different-length of the nsp7-10 region, BL21 Rosetta2 (Merck Millipore) were transformed, grown in culture flasks to 0.4-0.6 OD_600_ in 0.3-1 L, then induced with 50 μM anhydrotetracycline and continued to grow at 20°C overnight. To pellet, cultures were centrifuged (6000 x *g* for 20 min) and cells were frozen at −20°C. To separate soluble nsp’s, pelleted cells were lysed in 1:5 (v/v) buffer B1 (40 mM phosphate buffer, 300 mM NaCl) with one freeze-thaw cycle, sonicated (micro tip, 70 % power, 6 times on 10 s, off 60 s; Branson digital sonifier SFX 150) and then centrifuged (20,000 × *g* for 45 min). The proteins were isolated with Ni^2+^-NTA beads (Thermo Fisher Scientific) in gravity flow columns (BioRad). First the beads were equilibrated with 20 column volumes (CV) B1 + 20 mM imidazole, then bound to nsp’s by incubation with crude extract for 60 min and finally to remove unspecifically bound proteins washed with 20 CV B1 +20 mM imidazole followed by 10 CV of B1 +50 mM imidazole. To elute the nsp’s, eight fractions of 0.5 CV B1 +300 mM imidazole were collected. Immediately after elution, the fractions were supplemented with 4 mM DTT. For quality analysis, SDS-PAGE was performed.

### 1.5.3. FRET peptide assays

For SARS-CoV M^pro^ activity assays, Förster resonance energy transfer (FRET) peptide substrates (FPS) were commercially purchased (Eurogentec), designed as SARS-CoV cleavage site analogues of twelve amino-acid residues (P6-P6’) with a FRET pair labelling, namely the fluorophore (F) HiLyte488 at the N-terminus and the Quencher (Q) QXL520 at the C-terminus (Table S 1).

Initially, the freeze-dried FPS were dissolved in DMSO to 2 mM and stored at −20°C, protected from light. For sample preparation, the peptides were serially diluted in 50 mM HEPES, 10% (v/v) glycerol, 1 mM DTT, pH 7.5 and final concentrations were adjusted by measuring A_280_ from five independent droplets, using ε = 73,000 mol^−1^ cm^−1^, the absorption coefficient of the HiLyte488 fluorophore as given by the manufacturer.

Assays were carried out using a 96-well plate reader (Infinite200, Tecan) with the following parameters: excitation wavelength 485 nm (bandwidth 9 nm), emission wavelength 535 nm (bandwidth 20 nm), gain 80, 10 flashes and 40 μs integration time. Further, flat bottom black polystyrol 96-well plates (Greiner) were used for fluorescence applications.

Fluorescence coefficients k_sx_ and inner filter effect of the peptides were determined for each substrate according to previous protocols [32] (Figure S 2). The kinetic parameters were determined as suggested by Grum-Tokars et al. [25]. Briefly, 20 μL SARS-CoV M^pro^ (final conc. c_Mpro_ of 0.5 μM) was mixed with 80 μL FRET peptide substrates of six different concentrations (final conc. c_X,S_ of 2 μM, 1μM, 0.5 μM, 0.25 μM, 0.125 μM and 0 μM) and fluorescence (AFU) was monitored every 30 seconds for 5 minutes. To calculate kinetic parameters, triplicate measurements were performed.

The fluorescence coefficients k_sx_ were used to convert the initial increase of AFU into the initial reaction rate, which was then normalized by c_Mpro_ to rate/[Enzyme]. For comparing substrate specificities, the rates/[Enzyme] were plotted against the substrate concentrations c_X,S_ resulting in process curves. The slopes of the process curves correspond to the apparent k_cat_/K_M_.

The influence of salt conditions on enzyme activity was determined by incubating 20 μL M^pro^ (final concentration 0.1 μM) with 80 μM FRET peptide substrate FPS4-5 (final concentration 1 μM) in varying NH_4_OAc (10% Glycerol, pH7.5 and 50 mM, 165 mM, 280 mM, 395 mM, 510 mM, 625 mM, 740 mM, 855 mM, 970 mM, 1085 mM and 1200 mM) and NaCl (50 mM HEPES, 10% Glycerol, pH 7.5 and 0 mM, 120 mM, 240 mM, 360 mM, 480 mM, 600 mM, 720 mM, 840 mM, 960 mM, 1080 mM and 1200 mM). The fluorescence was followed every 60 seconds by monitoring (AFU), which initial slopes (AFU/s) were converted by the fluorescence coefficient k_sx_ into the initial rate (Figure S 1).

### 1.5.4. SDS PAGE

SDS-PAGE was performed with a 4-12% gradient acrylamide Bis-tris gel with XT MES running buffer. For *in-vitro* processing, 3.2 μM SARS-CoV M^pro^ was incubated with 13 μM SARS-CoV nsp7-9-His (ratio ~1:4) at 4°C in 20 mM phosphate buffer, 150 mM NaCl, 1 mM DTT at pH 8.0.

### 1.5.5. Native MS

To prepare samples for native MS measurements, they were buffer-exchanged into nano electrospray ionization (nanoESI)-compatible solution. Proteases M^pro^ were buffer exchanged into 250-500 mM NH_4_OAc, 1 mM pH 8.0 by two cycles of centrifugal gel filtration (Biospin mini columns, 6,000 MWCO, Biorad). The nsp’s were buffer-exchanged into 250-500 mM NH_4_OAc, 1 mM DTT, pH 8.0 by five rounds of dilution and concentration in centrifugal filter units (Amicon, 10,000 MWCO, Merck Millipore).

NanoESI capillaries were pulled in-house from borosilicate capillaries (1.2 mm outer diameter, 0.68 mm inner diameter, filament, World Precision Instruments) with a micropipette puller (P-1000, Sutter Instruments) using a squared box filament (2.5 × 2.5 mm, Sutter Instruments) in a two-step program. Subsequently, capillaries were gold-coated using a sputter coater (Q150R, Quorum Technologies) with 40 mA, 200 s, tooling factor 2.3 and end bleed vacuum of 8×10^−2^ mbar argon.

Native MS was performed at an nano-electrospray quadrupole time-of-flight (nanoESI-Q-TOF) instrument (Q-TOF2, Micromass/Waters, MS Vision) modified for higher masses [33]. Samples were ionized in positive ion mode with voltages applied at the capillary of 1300-1500 V and at the cone of 130-135 V. The pressure in the source region was kept at 10 mbar throughout all native MS experiments. For purpose of desolvation and dissociation, the pressure in the collision cell was adjusted to 1.3-1.5×10^−2^ mbar argon. Native mass spectra were obtained at accelerating voltage of 10-30 V while for CID-MS/MS these voltages were increased to 30-200 V. In ESI-MS overview spectra for nsp’s, the quadrupole profile was 1-10,000 *m/z*. In tandem MS, for precursor selection, LMres and HMres were adjusted at 10-30 V collisional voltage until a single peak was recorded, and then dissociation was induced.

To calibrate raw data, CsI (25 mg/ml) spectra were acquired and calibration was carried out with MassLynx (Waters) software. Data were analysed using MassLynx (Waters), Massign (by Nina Morgner [34]). All determined masses are provided (Table S 3).

To start the processing reactions, nsp’s and M^pro^ were incubated (ratio ~1:4–1:10). Three independent reactions were started in parallel and incubated at 4°C. To acquire mass spectra at defined time-points, sample aliquots of 1-3 μL were taken by means of a microliter syringe (5 μL, Hamilton) with flexible fused silica tubing (Optronis) and loaded into in-house fabricated nanoESI capillaries, which were mounted on the nanoESI source, all within two minutes. Then, raw spectra were acquired in the first 300 scans (1 scan/sec).

To analyze the data, the raw spectra were smoothed (2×5) in Masslynx 4.1 (Waters) and then nsp’s were assigned to peak series. For each assigned mass species, signal responses (SRs), using the combined relative intensities of peaks, were summarized and normalized. This was done independently for each spectrum. Finally, the average and standard deviation for SR and time-points was calculated from three independent spectra. Poorly resolved spectra were not included in the data analysis.

To determine mass fractions (MF), the SRs of assigned peaks for monomers and complexes were combined (Table S 2).

## 1.6. Acknowledgement

The Heinrich Pette Institute, Leibniz Institute for Experimental Virology is supported by the Free and Hanseatic City of Hamburg and the Federal Ministry of Health. C.U. and B.K. are supported by EU Horizon 2020 ERC StG-2017 759661. C.U. and B.K.․ also received funding through Leibniz Association through SAW-2014-HPI-4 grant. SF and LR thank the German Federal Ministry for Education and Research (BMBF) for funding (grants 01KX0806, 01KX0807, and 05K18FLA). RH acknowledges funding from the SILVER project of the European Commission (contract HEALTH-F3-2010-260644).

## 1.7. Supplement

- Supplementary figures and tables
- Author contributions
- Supplementary references

**Table S 1:** FRET peptide substrates (FPS) purchased to determine kinetic parameters of Mpro.

| SARS-CoV FPS | Product |
| --- | --- |
| F-nsp4-5(N-term) | H-C(HiLyte™ 488)TS AVLQ-OH (7aa) |
| nsp4-5(C-term)-Q | H-SGFRK(QXL® 520)-NH2 (5aa) |
| F-nsp4-5-Q | H-C(HiLyte™ 488)TSAVLQSGFRK(QXL®520)-NH2 (12aa) |
| F-nsp7-8-Q | H-C(HiLyte™ 488)NRATLQAIASK(QXL®520)NH2 (12aa) |
| F-nsp8-9-Q | H-C(HiLyte™ 488)SAVKLQNNELK(QXL®520)-NH2 (12aa) |
| F-nsp9-10-Q | H-C(HiLyte™ 488)ATVRLQAGNAK(QXL®520)-NH2 (12aa) |

**Table S 2:** Formulas to determine mass fractions:

|  |
| --- |
| MF SARS nsp7 = SR [nsp7 mon. + 2*nsp7 dim. + nsp7+8(1:1) + 2*nsp7+8(2:2)] |
| MF SARS nsp8 = SR [nsp8 mon. + nsp7+8(1:1) + 2*nsp7+8(2:2)] |
| MF SARS nsp9 = SR [nsp9 mon. + 2xnsp9 dim.] |
| MF SARS nsp10 = SR [nsp10 mon.] |
| MF 229e nsp7 = SR [nsp7 + nsp7+nsp7-9] |
| MF 229e nsp7-9 = SR [nsp7-9 + nsp7+nsp7-9] |

