## Supplementary material for "Processing of the SARS-CoV pp1a/ab nsp7-10 region": figures and tables

### 1. Supplementary

Affiliation: [1]Heinrich Pette Institute, Leibniz Institute for Experimental Virology, Martinistraße 52, 20251 Hamburg, Germany; [2]University of Hamburg, Institut für Biochemie und Molekularbiologie, Martin-Luther-King-Platz 6, 20146 Hamburg, Germany; [3]University of Lübeck, Institute of Biochemistry, Center for Structural and Cell Biology in Medicine, Ratzeburger Allee 160, 23562 Lübeck, Germany; [4]German Center of Infection Research (DZIF), Hamburg - Lübeck - Borstel - Riems Site, University of Lübeck, 23562 Lübeck, Germany [5]Deutsches Elektronen Synchrotron (DESY), Notkestraße 85, 22607 Hamburg, Germany; [6]European XFEL GmbH, Holzkoppel 4, 22869 Schenefeld, Germany.

#### 1.1. Supplementary figures and tables

##### 1.1.1. Molecular masses

Table S 1: Molecular weights (MWs) of nsp's, their cleavage products and complexes. Average theoretical mass (MWTheo.) according to amino acid sequence (Protein ID: P0C6X7). Molecular weight (MW) (MWExp.) determined from Native MS or CID (*) and listed with standard deviations (SD) (N=3) and the average full-width half maximum (FWHM) (N=3) of the main peak. All values in Da. The masses are given without SD for mass species that were safely assigned in only two spectra.

| Protein | $MW_{Theo}$ | $MW_{Exp.}$ | SD | FWHM |
|---|---|---|---|---|
| **Processing (SARS-CoV His-nsp7-10)** | | | | |
| His-nsp7 | 10677 | 10680.1 | 0.2 | 1.4 |
| nsp8 | 21866 | 21871 | 1 | 1.4 |
| nsp9 | 12401 | 12403.5 | 0.2 | 1.3 |
| nsp10 | 14843 | 14974 | 1 | 1.9 |
| His-nsp7-8 | 32525 | 32541 | 11 | 1.4 |
| His-nsp7-9 | 44908 | 44937 | 15 | 3.9 |
| His-nsp7-10 | 59734 | 59930 | 40 | 14.4 |
| **Complex formation (from nsp7-9-His)** | | | | |
| nsp7 | 9267.8 | 9268.3 | 0.1 | 2.4 |
| nsp8 | 21866.0 | 21867.3 | 0.5 | 3.7 |
| nsp8(2) dim | 43713.9 | 43700* | 30 | 16 |
| nsp7+8(2:1) | 40365.6 | 40404* | 8 | 3.6 |
| nsp7+8(1:2) | 52963.7 | 53009* | 6 | 4.5 |
| nsp7+8(2:2) | 62213.5 | 62300 | 10 | 7.1 |

##### 1.1.2. Specific substrate efficiencies from FRET peptide assay

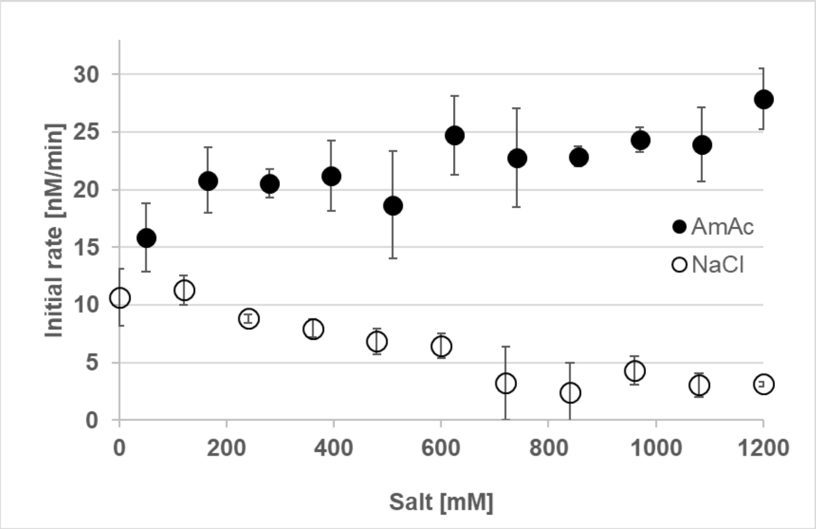


Figure S 1: Influence of $NH_4$Oac and NaCl on $M^{pro}$ initial rate. While activity of SARS-CoV $M^{pro}$ was slightly increasing with $NH_4$OAc concentration, it was inhibited by increasing NaCl concentration.

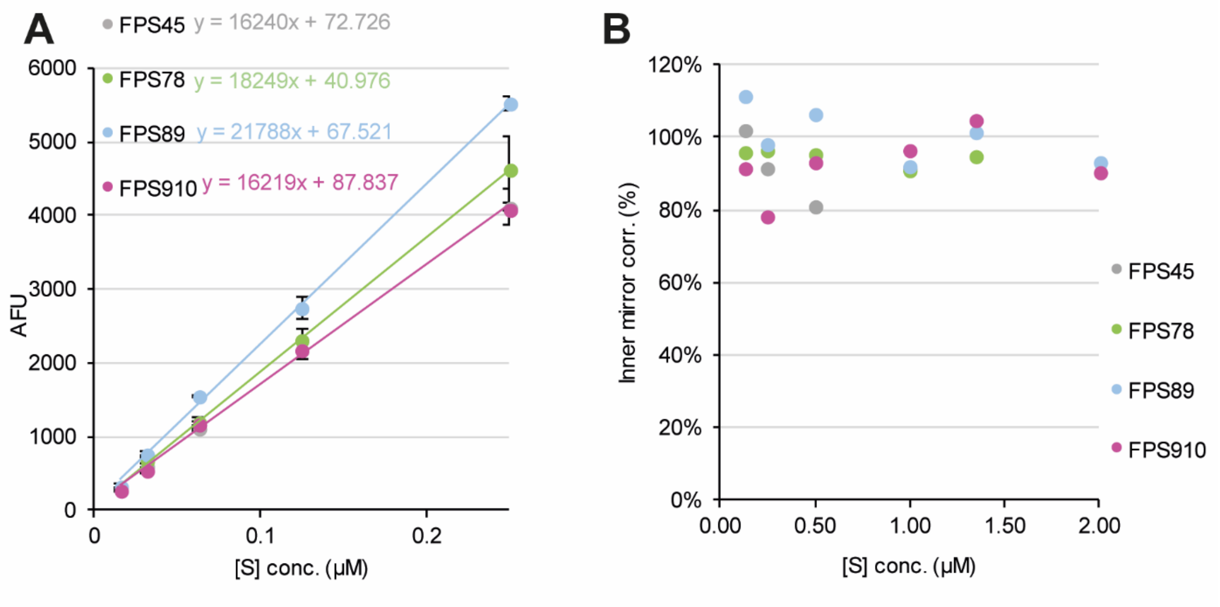


Figure S 2: Determination of specific fluorescence coefficient $k_{sx}$ and inner filter effect. **(A)** Specific FPS fluorescence determined from the slope of arbitrary fluorescence units at different concentrations of completely cleaved substrate (conc.[S]). The found specific fluorescence was used to relate measured AFU increase in activity assays to concentration of cleaved peptide. **(B)** Inner filter correction for varying substrate concentrations (conc.[S]). Since no clear trend was apparent the inner filter effect appears not to play a critical role in the used experimental setup and the correction was not applied for data analysis of other measurements.

##### 1.1.1. Processing of nsp7-10

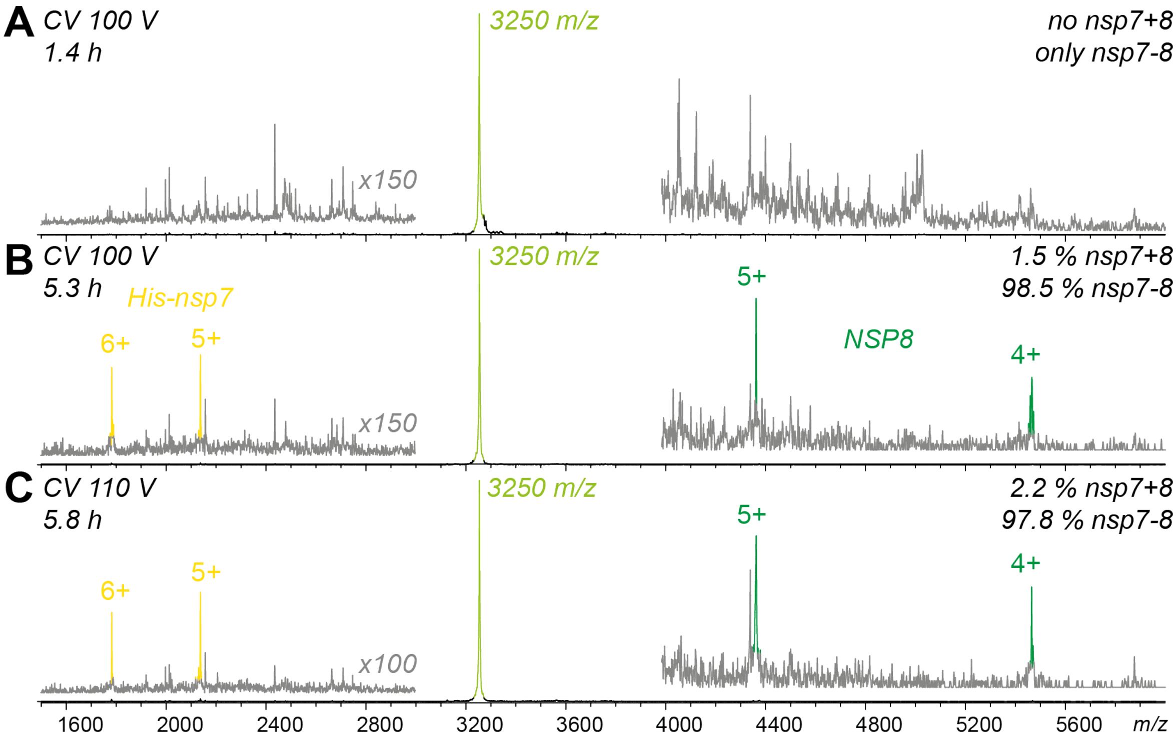


Figure S 3:SARS nsp7-10 processing in native MS: CID of nsp interactions. Product ion spectra (black) of His-nsp7-8/nsp His-7+8 precursor (3250 *m/z*, 10+, light green) at three time-points of nsp7-10 processing (ESI-MS spectra of processing). Superimposed product ion spectra (grey) magnified at indicated ratio. The ratio of His-nsp7-8 to nsp His-7+8 (1:1) is given, based on the ratio of intensity of precursor ion peak to nsp7 product ion peaks for (A) 1.4 h, (B) 5.3 h, and (C) 5.8 h. After 5.8 h, when the precursor ion is the dominant peak in the ESI-TOF spectrum, only 2.2 % of the molecular ion is dimeric, and 97.8 % is covalently bound.

Table S 2Substrates and their relative efficiency for $M^{pro}$. Given is the name of the substrate, the amino-acid sequence P6'-P6 at the nsp inter-domain junction cleavage site, relative efficiency, determined apparent efficiency ($k_{cat}/K_M$) as determined in FRET peptide protease assay. Highlighted (blue) is a conserved NNE motif of the non-canonical nsp8-9 cleavage site. Apparent $k_{cat}/K_M$ of $M^{pro}$ (0.5 µM) with different concentrations of FRET peptide substrates (FPS) in 50 mM HEPES, 10 % Glycerol pH7.5.

| Name | Amino acids | Rel. efficiency (%) | Apparent $k_{cat}/K_M$ ($\mu M^{-1}$ $min^{-1}$) ±SD , (N=3) |
|---|---|---|---|
| FPS4-5 | TSAVLQ↑SGFRK | 100 | 0.106±0.005 |
| FPS7-8 | NRATLQ↑AIASK | 17.1 | 0.018±0.003 |
| FPS8-9 | SAVKLQ↑NNELK | 0.1 | 0.0001±2x$10^{-5}$ |
| FPS9-10 | ATVRLQ↑AGNAK | 51.8 | 0.0547±0.002 |

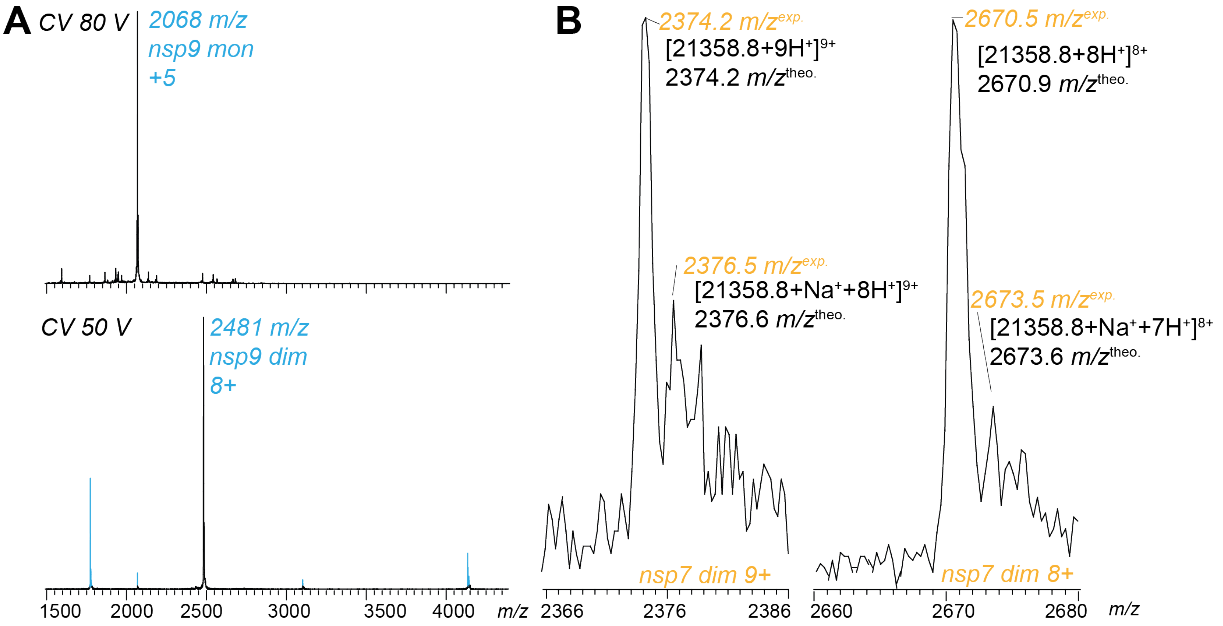


Figure S 4:Homo-dimers of nsp7 and nsp9 identified in SARS-CoV His-nsp7-10 processing. (A) Collision-induced dissociation to identify ions of monomeric and dimeric mass species of nsp9. Product ion spectra of 2068 *m/z* (top) precursor shows onset of backbone fragmentation at 80 V CV, while 2481 *m/z* precursor (bottom) shows dissociation at 50 V CV. (B) Zoom of two charge states of a peak series assigned to nsp7 dimer in ESI-MS spectra of processing. Sodium adducts of experimental $m/z^{exp}$ are well in line with theoretical $m/z^{theo}$ that are expected from a nsp7 dimer. A monomeric nsp7 could not have adducts at the indicated $m/z^{exp}$. Affiliated native MS spectra in main part (Figure 2).

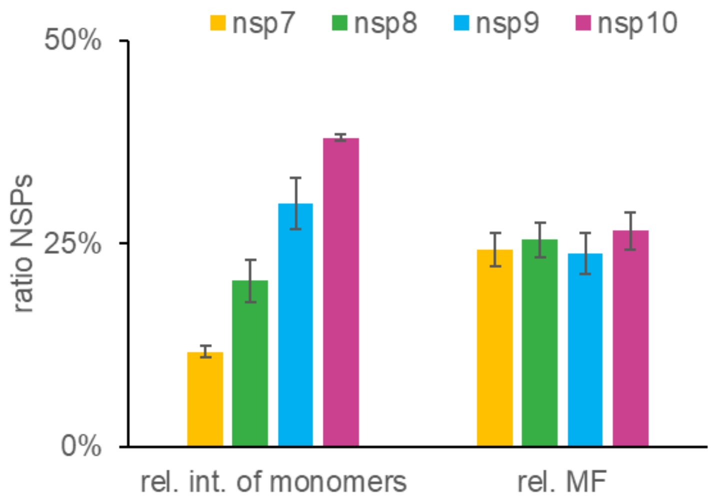


Figure S 5:Signal response ratio of SARS-CoV nsp7-10 domains.Compared to the monomeric mass species the signals converted to mass fractions of nsp7, nsp8, nsp9 and nsp10 were found evenly distributed, indicating that they well represent the equimolar presence of nsp's (Equations listed in Table S 2). This result shows that a plot of the mass fractions could be used to observe relative concentrations directly from the ratios. Furthermore, instrument parameters and response factors appear to have less influence than expected. However, previously it was shown that with a similar instrument, larger species give more intense signals [1]. Also here, the tested nsp's (~9.5-15 kDa) may have different ion efficiencies than larger species, e.g. nsp7-10 and $M_{pro}$ (~59 kDa and ~69 kDa), and should therefore not be compared directly. The rel. int. ratio of monomers by using the intensities of all charge states assigned to monomers (AVG.±SD, N=3; 10±1%, 19±3%, 30±4% and 41±1%). The ratio of mass fraction (MF) is the SR ratio, corrected for all non-covalent complexes (AVG.±SD, N=3; 23±3%,

26±3%, 24±3% and 28±3%).

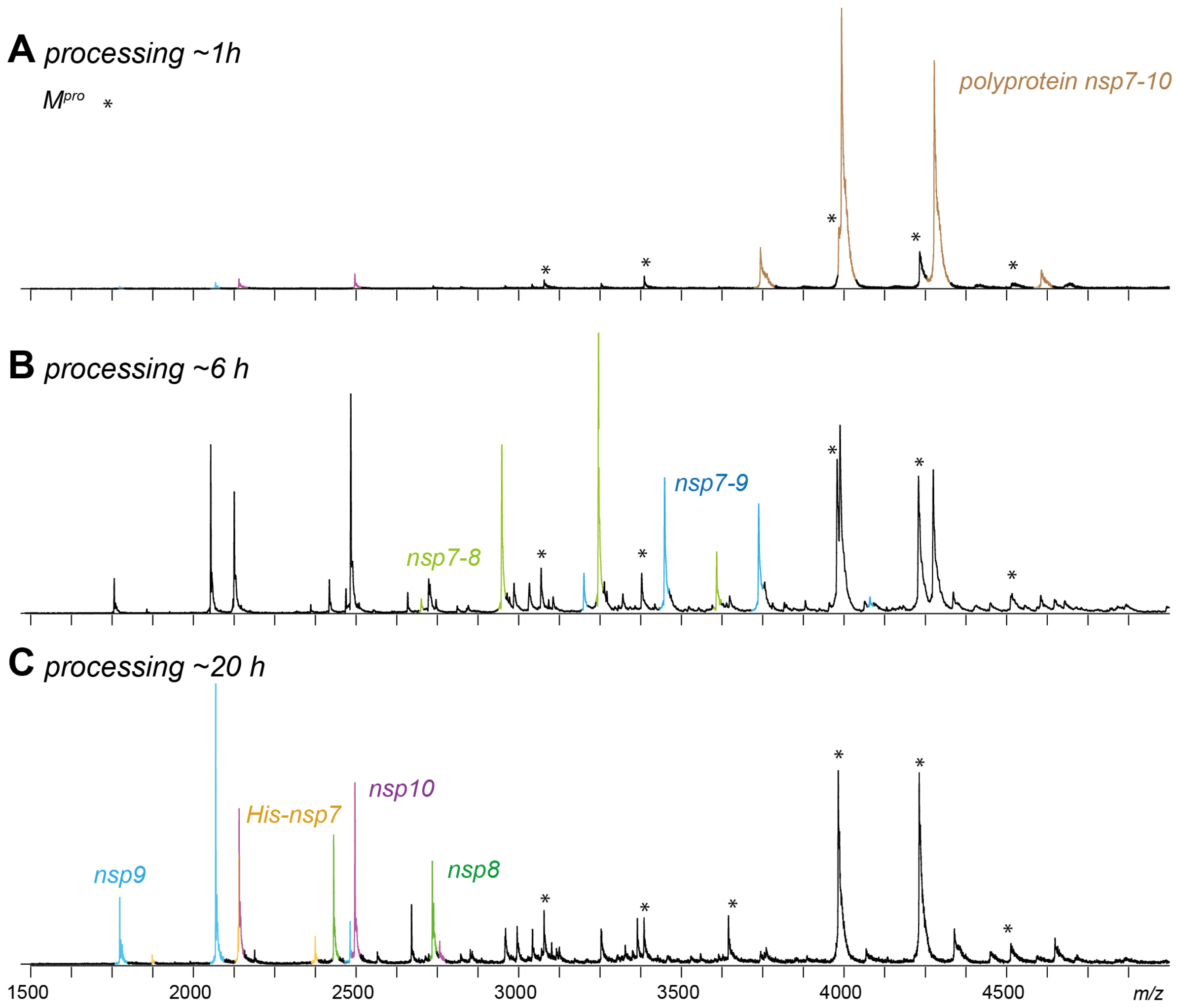


Figure S 6: SARS-CoV nsp7-10 processing in native MS: Exemplary spectra. Exemplary native mass spectra illustrate processing from polyprotein to nsp end-products. For *in-vitro* processing, 1.25 µM SARS-CoV M$^{pro}$ and 12.5 µM SARS-CoV His-nsp7-10 (~1:10 ratio) were incubated at 4°C in 250 mM $NH_4OAc$, 1 mM DTT at pH 8.0. After mixing the components, samples were injected into an electrospray capillary and native MS spectra were recorded at indicated time points. (A) Mass spectrum at 1 h showing nsp7-10 (beige). Additionally indicated are peak signals of M$^{pro}$ monomer and dimer (asterisk), as well as low-intensity signals of nsp9 (blue) and nsp10 (pink). (B) Mass spectrum at 6 h with dominant peaks assigned to intermediate products nsp7-8 and nsp7-9. (C) Mass spectrum at 20 h showing dominant peaks assigned to mature cleavage products His-nsp7 (yellow), nsp8 (green), nsp9 and nsp10.

#### Processing of nsp7-9-His

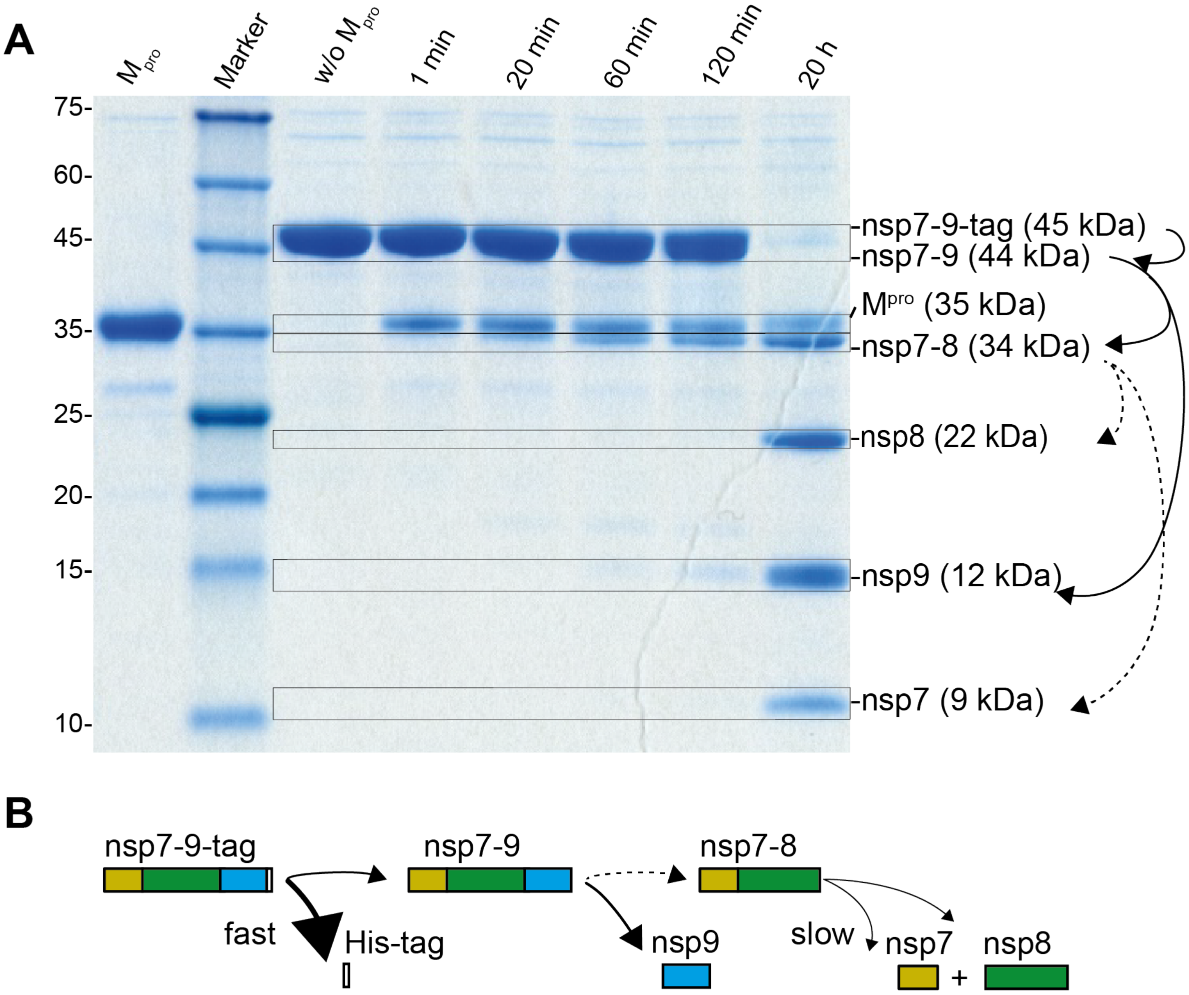


Figure S 7: SDS-PAGE of SARS nsp7-9-His processing. To start the reaction, 3.2 µM SARS-CoV M$^{pro}$ was incubated with 13 µM SARS-CoV nsp7-9-His (ratio ~1:4) at 4°C in 20 mM phosphate buffer, 150 mM NaCl, 1 mM DTT at pH 8.0. (A) Lane 1, SARS-CoV M$^{pro}$; lane 2, Roti Tricolor marker; lane 3, nsp7-9-His; lane 4-8, *in-vitro* processing of nsp7-9 with M$^{pro}$ after indicated time points. Frames indicate for band assignment. Black arrows illustrate the cleavage pathway as concluded from band shift. (B) Symbols illustrate sequential cleavage of nsp mass species, as concluded from the nsp7-9-His processing experiments.

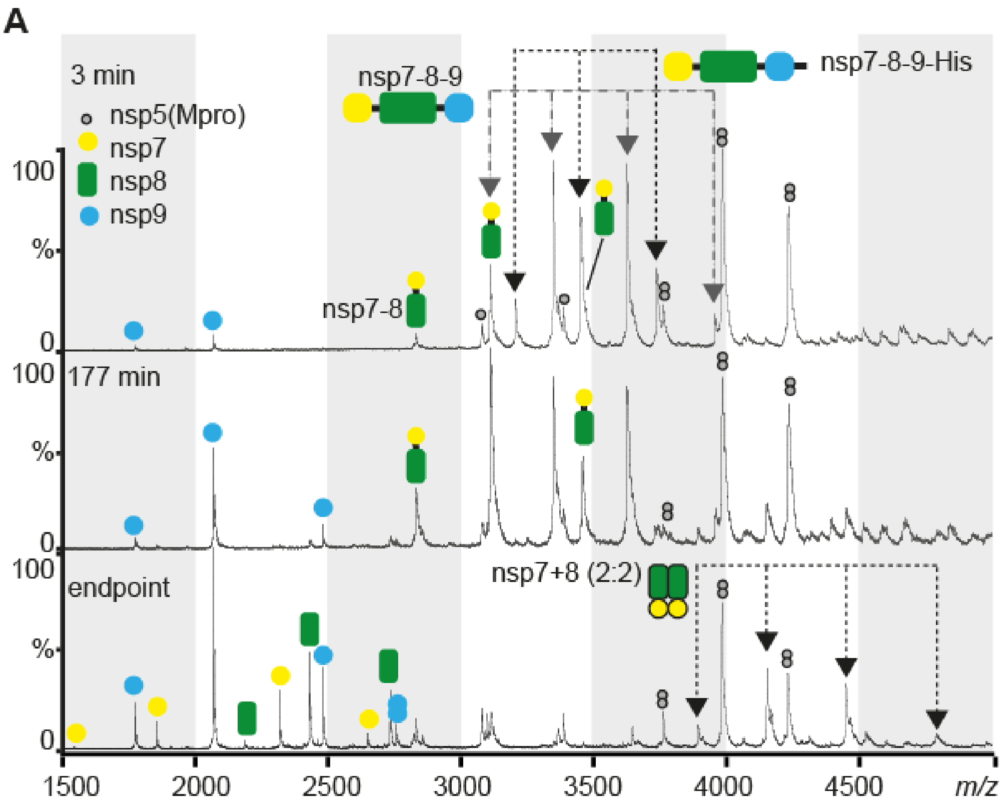


Figure S 8: Native MS exemplary spectra of SARS-CoV nsp7-9-His processing. Exemplary native MS spectra showing three time-points of sampling. For in *in-vitro* processing, 2 µM M$^{pro}$ was incubated with 14 µM nsp7-9 (ratio 1:7) at 4°C in 250 mM $NH_4OAC$, 1 mM DTT, pH 8.0. Top: Native mass spectrum after 3 min. Middle: Native mass spectrum after 177 min. Bottom: Native mass spectrum after 20 h.

##### 1.1.1. Complex formation of nsp7+8

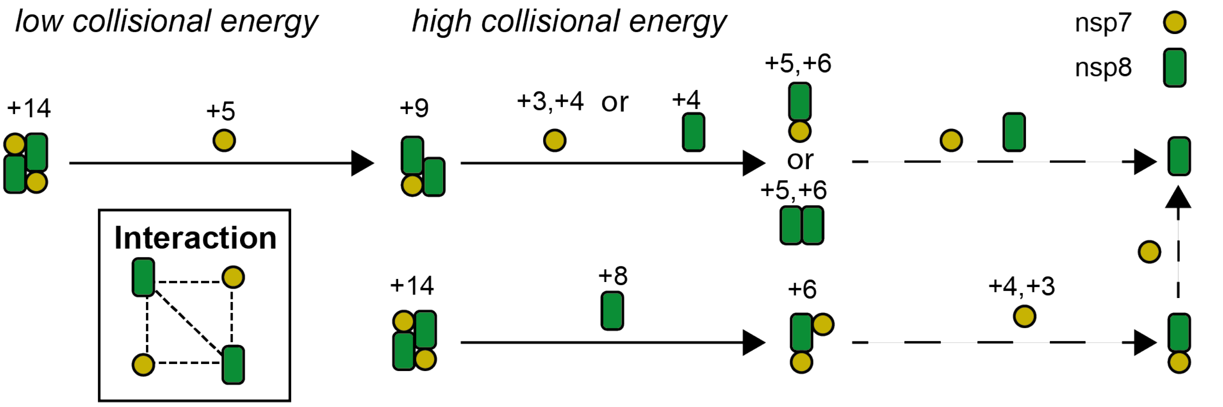


Figure S 9: Gas-phase dissociation pathways of nsp7+8(2:2). Two alternative dissociation pathways of SARS-CoV tetramer and interaction of subunits within the tetramer as concluded from CID spectra. Main pathway (top): Initially, nsp7+8(2:2) dissociates into nsp7 and nsp7+8(1:2), suggesting a peripheral positioning of nsp7 within the complex. In a follow-up dissociation, nsp7+8(1:1) is detected at elevated collisional energy. Alternative pathway (bottom): At higher collisional energy, dissociation into nsp8 and nsp7+8(2:1) is preferred. nsp8 dimers appear as well. Interaction map: Subunit interactions as concluded from CID-MS results. Symbols illustrate molecular ions from different mass species found as dissociating ions in CID. Black arrows indicate for the ejected ions. Dashed arrows indicate for unobserved dissociations that most likely follow up. Charge states labelled taken from dissociating ions found in CID of SARS-CoV nsp7+8 (2:2) (+14) and when combined match the charge state of the precursor ion.

Table S 3: Buried surface areas as an approximation for binding interface. To get an approximation of buried interface, the surface area of single subunits was subtracted by the surface area of T1 or T2 with PyMol and get_area input order. Here, nsp7/nsp8 in T1/2 is the buried surface between a single subunit and the complex. nsp8I:nsp8I/nsp8II:nsp8II is the buried surface by the nsp8 scaffold alone.

| Interface | Buried surface (Å$^2$) | Buried surface (%) |
|---|---|---|
| T1 total | 10281 | 100 |
| nsp7 in T1 | 2153 | 21 |
| nsp8I in T1 | 2988 | 29 |
| nsp8I:nsp8I | 835 | 8 |
| T2 total | 10377 | 100 |
| nsp7 in T2 | 1988 | 19 |
| nsp8II in T2 | 3201 | 31 |
| nsp8II:nsp8II | 1214 | 12 |

#### 1.1. Author contributions

Conceptualization and methodology B.K and C.U.; Cloning constructs S.F.; providing research materials R.H.; investigation B.K., discussion of results B.K., C.U,, S.F. and L.R.; formal analysis, visualization and original draft B.K and C.U; writing - review and editing B.K., C.U., S.F., L.R., and R.H..
